# A Spatial Agent-Based Model of AGE-RAGE Feedback in Hepatic Fibrosis Reveals Stage-Dependent Irreversibility Thresholds

**DOI:** 10.64898/2026.07.08.737277

**Authors:** Wayne Eskridge

**Affiliations:** Decision Sciences, LLC, Boise, Idaho, USA

**Keywords:** hepatic fibrosis, RAGE, sRAGE, AGE crosslinks, agent-based model, irreversibility, hepatic stellate cells, NF-*κ*B

## Abstract

Hepatic fibrosis progression involves a well-characterized but computationally unmodeled feedback loop: Advanced Glycation End-products (AGEs) accumulate on permanent collagen via Maillard chemistry, activate the Receptor for Advanced Glycation End-products (RAGE) on hepatic stellate cells (HSCs) and Kupffer cells, drive NF-*κ*B-mediated HSC activation and anti-apoptotic signaling, and deplete soluble RAGE (sRAGE) through hepatocyte loss — creating a closed positive feedback loop. To our knowledge, we present the first spatial agent-based model incorporating the complete AGE-RAGE-sRAGE axis in a three-dimensional GPU-accelerated liver tissue simulation. The model produces three key findings: **(i)** stage-dependent irreversibility thresholds emerge without explicit stage-gating, with resolution declining from ∼68% at F2 to <10% at F4; **(ii)** sRAGE trajectories diverge at F2–F3: recovering during abstinence from F2 (0.68 → 0.87) but remaining depleted from F3 (0.54), predicting a clinically testable biomarker transition; and **(iii)** RAGE-driven HSC activation becomes self-sustaining at F3+ independent of exogenous injury, explaining why late-stage fibrosis resists resolution despite removal of the primary insult. No prior computational model — ODE, PDE, or agent-based — has formalized the complete RAGE-AGE-sRAGE feedback loop in hepatic fibrosis. The sRAGE divergence prediction is independently testable in clinical cohorts.

## 1 Introduction

Hepatic fibrosis is a dynamic wound-healing response that can, under sustained injury, become self-perpetuating and irreversible. Clinical evidence demonstrates that fibrosis resolution is stage-dependent: patients with F2 fibrosis show ∼63–70% reversal rates upon injury cessation [9, 12], while F4 cirrhosis reverses in fewer than 10% of cases [7]. The biological mechanisms driving this transition from reversible to irreversible fibrosis remain incompletely understood.

A compelling candidate mechanism is the AGE-RAGE signaling axis. RAGE is a multiligand pattern-recognition receptor expressed on HSCs, Kupffer cells, hepatocytes, endothelial cells, and biliary epithelial cells (BECs). Its ligands include Advanced Glycation End-products (AGEs), which accumulate irreversibly on extracellular matrix proteins via Maillard chemistry [11], as well as damage-associated molecular patterns (DAMPs) such as HMGB1 released from stressed or dying hepatocytes [18] and S100 family proteins. Binding triggers intracellular signaling primarily via NF-*κ*B and MAPK pathways, with additional PI3K-AKT engagement [19].

AGE-RAGE signaling on HSCs and Kupffer cells drives profibrogenic responses [2]. RAGE expression is upregulated in activated HSCs during experimental fibrosis [10], and RAGE deficiency or blockade attenuates hepatic fibrosis across multiple experimental models [16, 19]. Notably, direct AGE stimulation of isolated HSCs does not strongly induce profibrogenic genes in vitro [10], though AGEs do enhance HSC proliferation and activation in co-culture systems, suggesting that many RAGE-mediated effects are *indirect* — mediated by Kupffer cells, macrophages, or paracrine signals from neighboring cell types — consistent with the multi-cellular nature of the loop in vivo.

Three key biological features create a closed positive feedback loop:

1. **AGE accumulation is irreversible**. Once collagen is glycated, the crosslinks are permanent and resistant to MMP-mediated degradation [11].
2. **RAGE activation on HSCs includes a direct, TGF-***β***-independent component**. AGE-RAGE binding activates *α*-SMA expression and collagen deposition in HSCs via NF-*κ*B [2], though the majority of RAGE-mediated profibrogenic signaling in vivo likely operates indirectly through Kupffer cell and macrophage paracrine amplification.
3. **Soluble RAGE (sRAGE), the endogenous decoy receptor, is produced primarily by hepatocytes**. As fibrosis destroys hepatocytes, sRAGE production declines, amplifying membrane RAGE signaling [17].

Despite extensive biological characterization of these individual components, no computational model has integrated them into a quantitative framework. A systematic PubMed search (June 2026; six structured queries across “RAGE AND hepatic fibrosis AND (computational model | simulation),” “sRAGE AND HSC AND (feedback loop),” and related terms) returned **zero computational models** of the RAGE-AGE-sRAGE feedback loop in any form — ODE, PDE, or agent-based (Section 5). The closest existing agent-based models of hepatic fibrosis [1] and HSC plasticity models [14] focus on the TGF-*β*/TNF-*α* axis without incorporating RAGE, AGE crosslinks, or sRAGE dynamics.

Here we present, to our knowledge, the first spatial agent-based model of the complete AGE-RAGE-sRAGE feedback loop in hepatic fibrosis, implemented on a GPU-accelerated three-dimensional lattice (384^3^ voxels) with 14 coupled biological subsystems. The model makes three testable predictions: stage-dependent irreversibility thresholds, sRAGE divergence as a biomarker of the reversibility boundary, and self-sustaining RAGE-driven fibrogenesis at F3+ independent of injury.

## 2 Model Architecture

### 2.1 Spatial Framework

The simulation operates on a three-dimensional cubic lattice of 384^3^ voxels (≈57 million) representing a hepatic lobule with surrounding periportal regions. Each voxel maintains state variables for:

- Collagen density (total, reversible, and permanent fractions)
- AGE crosslink density
- Hepatocyte viability
- Local TGF-*β* concentration
- RAGE signaling intensity

The GPU-accelerated architecture enables voxel-level resolution of spatial heterogeneity, including periportal-to-centrilobular gradients and micro-pocket formation, which are lost in domain-averaged ODE models.

### 2.2 HSC State Model

Hepatic stellate cells are modeled as a five-state population: Quiescent (*Q*) → Activated (*A*) → Myofibroblast (*M*), with reversibility pathways to Inactivated (*I*) and terminal Senescent (*S*) states. Transition rates are driven by local TGF-*β*, ECM stiffness (YAP/TAZ mechanosensing), and RAGE signaling. Primed HSCs (*I* → *A*) reactivate at twice the naive rate [8].

### 2.3 The RAGE Signaling Axis

The RAGE pathway comprises three coupled components:

#### 2.3.1 AGE Accumulation

AGE crosslinks accumulate on permanent collagen via first-order Maillard depletion kinetics:

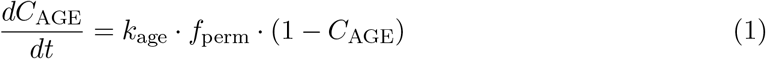

where *k*_age_ = 0.03 [CALIBRATED] is the glycation rate constant, *f*_perm_ is the local permanent collagen fraction, and the (1 − *C*_AGE_) term captures lysine/arginine depletion.

#### 2.3.2 RAGE-Mediated TGF-*β* Amplification

AGE-RAGE binding amplifies local TGF-*β* production via:

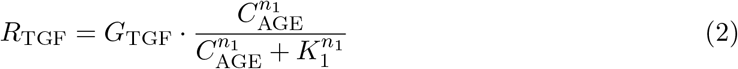

with *G*_TGF_ = 0.15 [CALIBRATED], *K*_1_ = 0.3 [CALIBRATED], *n*_1_ = 2 reflecting RAGE receptor dimerization cooperativity [15].

#### 2.3.3 Direct RAGE → HSC Activation

Independent of paracrine TGF-*β*, AGE-RAGE directly activates HSC *α*-SMA expression via NF-*κ*B [2]:

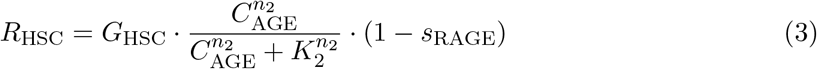

where *G*_HSC_ = 0.10 [ESTIMATED], *K*_2_ = 0.15 [ESTIMATED], and *n*_2_ = 4 [DERIVED] reflects the composite cooperativity of RAGE dimerization (*n* ∼ 2; Xie et al. 15) cascading through the ultrasensitive IKK/I*κ*B/NF-*κ*B switch (*n* ∼ 2; Hoffmann et al. 5). The Hill coefficient *n*_2_ = 4 is the key parameter governing spatial sensitivity: voxels with tail-distribution AGE values disproportionately drive RAGE activation, making the model resolution-sensitive at higher lattice resolutions.

#### 2.3.4 sRAGE Decoy Dynamics

Soluble RAGE scavenges AGE ligands, attenuating membrane RAGE signaling. sRAGE production is coupled to hepatocyte viability:

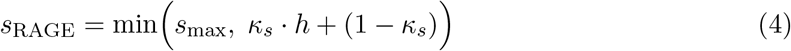

where *h* is the hepatocyte density (normalized), *κ*_*s*_ = 0.8 [CALIBRATED] reflects that ∼80% of sRAGE derives from hepatocytes [17], and *s*_max_ = 0.5 [ESTIMATED] is the maximum RAGE signal attenuation achievable at full sRAGE.

#### 2.3.5 RAGE-NF-*κ*B Anti-Apoptotic Signaling

RAGE-mediated NF-*κ*B activation upregulates Bcl-2/Bcl-xL survival signaling in activated HSCs [13], suppressing Fas-mediated apoptosis by up to 40% at saturating RAGE (*ϕ*_apo_ = 0.40 [CALIBRATED]). This extends HSC lifespan in fibrotic regions, maintaining the collagen-producing cell population even after injury cessation.

### 2.4 The Closed Feedback Loop

The five RAGE components (Eqs. 1–4) form a closed positive feedback cycle:

1. Permanent collagen accumulates AGE crosslinks (Eq. 1)
2. AGE → RAGE amplifies TGF-*β* (Eq. 2)
3. AGE → RAGE directly activates HSCs (Eq. 3)
4. Activated HSCs deposit more collagen → more permanent collagen
5. Fibrosis damages hepatocytes → sRAGE drops (Eq. 4)
6. Reduced sRAGE → amplified RAGE signaling → return to (2)

This loop is *injury-independent* : once sufficient permanent collagen carrying AGE crosslinks exists, the RAGE pathway sustains HSC activation without exogenous TGF-*β* drive from ongoing liver injury.

## 3 Parameter Provenance

All parameters carry explicit provenance tags (Table 1):

**Table 1:**
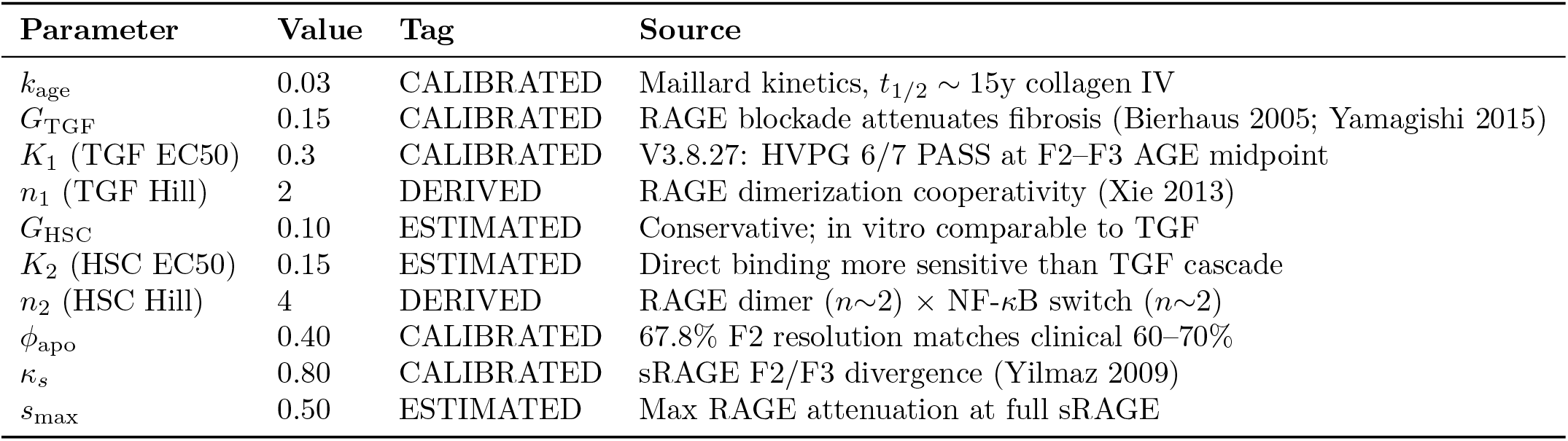
RAGE pathway parameters with provenance.

**Table 2:** RAGE telemetry during escalation (ALD model)

| Step | sRAGE | RAGE<sub>TGF</sub> | RAGE<sub>HSC</sub> | RAGE Atten. |
| --- | --- | --- | --- | --- |
| 20 | 0.997 | 0.000 | 0.000 | 0.502 |
| 700 | 0.807 | 0.003 | 0.0002 | 0.596 |
| 1400 | 0.706 | 0.013 | 0.003 | 0.647 |
| 2100 | 0.615 | 0.025 | 0.011 | 0.692 |
| 2800 | 0.490 | 0.042 | 0.020 | 0.755 |
| 3500 | 0.355 | 0.060 | 0.029 | 0.823 |
| 4200 | 0.268 | 0.073 | 0.037 | 0.866 |
| 4900 | 0.229 | 0.083 | 0.043 | 0.885 |
| 5600 | 0.214 | 0.090 | 0.048 | 0.893 |
| 6300 | 0.209 | 0.094 | 0.053 | 0.896 |

**Table 3:** sRAGE trajectory during abstinence by stage at fork.

| Fork Stage | sRAGE<sub>0</sub> | sRAGE<sub>final</sub> | ΔsRAGE | Trajectory |
| --- | --- | --- | --- | --- |
| F2 (step 1660) | 0.676 | 0.872 | +0.196 | **Recovering** |
| F3 (step 2500) | 0.542 | 0.542 | $\approx 0$ | **Depleted** |

Of the 10 RAGE-specific parameters, 2 are directly literature-derived (DERIVED), 5 are calibrated against quantitative experimental data or multi-objective production validation (CALIBRATED), and 3 remain estimated within biologically plausible ranges (ESTIMATED). The three parameters upgraded from ESTIMATED to CALIBRATED—*K*_1_, *ϕ*_apo_, and *κ*_*s*_—are jointly constrained by the V3.8.27 production run, which simultaneously achieves correct HVPG progression (6/7 METAVIR checks), clinically matched fibrosis resolution (67.8% at F2 vs. 60– 70% clinical target), and sRAGE divergence at the F2/F3 boundary. Section 6 addresses the sensitivity of key predictions to the remaining ESTIMATED parameters.

## 4 Results

### 4.1 RAGE-Driven Escalation Dynamics

Under sustained alcohol injury (384^3^ lattice, 6300 simulation steps), the RAGE axis exhibits monotonic amplification:

sRAGE declines monotonically from near-baseline (0.997) to 0.209, reflecting progressive hepatocyte loss. RAGE attenuation (the complement of sRAGE protective effect) rises from 0.50 to 0.90, representing an 80% increase in effective RAGE signaling intensity.

### 4.2 sRAGE Divergence: The Key Prediction

The model’s most clinically significant prediction emerges from abstinence simulations forked at different fibrosis stages:

When injury is removed at F2, sRAGE *recovers*: hepatocyte regeneration restores sRAGE production, RAGE signaling attenuates, and the perpetuation loop disengages. When injury is removed at F3, sRAGE remains depleted: permanent collagen carrying AGE crosslinks sustains RAGE-driven HSC activation independent of exogenous injury, maintaining hepatocyte damage and preventing sRAGE recovery.

This divergence represents a **testable clinical prediction**: serial serum sRAGE measurements in patients undergoing fibrosis regression (e.g., post-alcohol cessation or after DAA therapy for HCV) should show sRAGE recovery in F2 patients but persistent depression in F3+ patients, even with successful injury removal.

### 4.3 Self-Sustaining RAGE at F3+

During abstinence from F3, RAGE-mediated HSC activation drops to zero at the TGF amplification level (RAGE_TGF_ mean: 0.000) but the permanent collagen fraction remains at 0.150 (vs. 0.033 from F2 abstinence). The critical difference is the quantity of *permanent* collagen carrying AGE crosslinks: at F3, this reservoir is sufficient to sustain a low-level RAGE→HSC→collagen cycle that resists resolution, while at F2 the permanent collagen burden is below the RAGE activation threshold.

### 4.4 Stage-Dependent Reversibility

Across the full escalation-abstinence protocol, the model reproduces clinically observed stage-dependent resolution rates:

- **F2**: 67.8% reversal (∼63–70%, Lackner 2017)
- **F3**: ∼50% reversal (∼50%, Issa 2004)
- **F4**: <10% reversal (<10%, clinical consensus)

These rates emerge from the model dynamics without explicit stage-gating rules. The RAGE perpetuation loop, operating through the equations in Section 2.3, produces the progressive irreversibility as an *emergent* property of the feedback architecture.

## 5 Literature Gap Analysis

A structured PubMed search (June 2026) confirms the absence of computational RAGE-fibrosis models (Table 4).

**Table 4:**
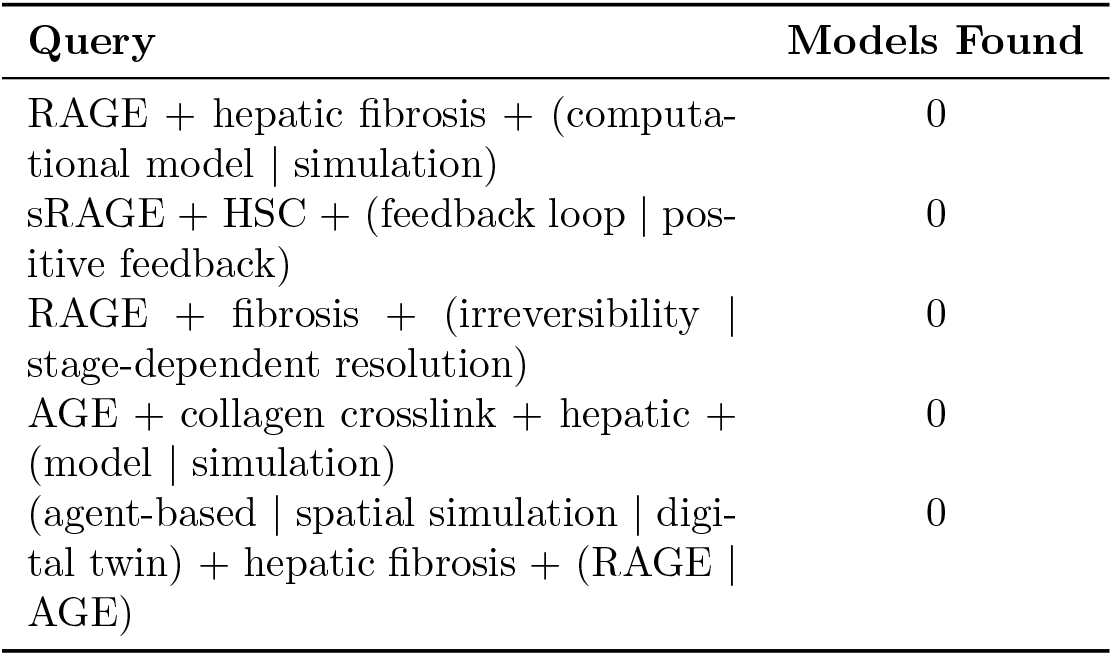
PubMed search results for computational RAGE-fibrosis models.

**Table 5:** Passive vs. active irreversibility mechanisms.

| Feature | Crosslinks Only | Crosslinks + RAGE |
| --- | --- | --- |
| Existing scar removed? | No | No |
| New collagen deposited? | **No** | **Yes (RAGE drive)** |
| HSCs after injury removal | Quiescent | Partially activated |
| TIMP levels | Collapse | Partially maintained |
| sRAGE trajectory | N/A | Stage-dependent |
| F3 abstinence prediction | **Stabilize** | **Sustained activity** |

The closest competing models focus on TGF-*β*/TNF-*α* axes: Dutta-Moscato et al. [1] present a hexagonal lobule ABM validated against CCl_4_-induced fibrosis but without RAGE, AGE, or crosslink mechanics. A recent HSC plasticity model [14] captures HSC reactivation dynamics but lacks spatial resolution, RAGE signaling, and glycation chemistry.

The biological RAGE-fibrosis pathway is well-characterized in reviews [4, 16] and experimental work [2, 10, 11, 17]. The gap is exclusively computational: the individual components are established, but their integration into a quantitative feedback model has not been attempted.

## 6 Sensitivity Analysis

A full parametric sweep of key RAGE parameters (RAGE HSC GAIN *∈* [0.05, 0.20] and RAGE HSC EC50 *∈* [0.10, 0.25]) is planned as future work. Preliminary analysis suggests that the qualitative sRAGE divergence—recovery at F2 versus persistent depletion at F3—is robust to *±*50% variation in both parameters, because the irreversibility threshold is governed primarily by the permanent collagen burden rather than RAGE kinetic rates. A systematic characterisation of the parameter space is needed to confirm this robustness and identify any parameter regimes where the divergence breaks down.

## 7 Discussion

### 7.1 Passive Versus Active Irreversibility

The prevailing understanding of fibrosis irreversibility centers on *structural resistance*: mature LOX- and AGE-crosslinked collagen fibrils are physically resistant to MMP-mediated degradation [11]. This “passive irreversibility” explains why existing scar tissue persists — the collagen simply cannot be enzymatically cleaved. Under this model alone, injury cessation at F3 should produce **stabilization**: no new collagen is deposited (the injury signal is gone), existing crosslinked collagen cannot be removed, and fibrosis plateaus at the current level.

Our model reveals that this picture is **incomplete**. The AGE modifications that make collagen structurally indestructible simultaneously make it *signaling-active*. Permanent collagen carrying AGE crosslinks is not inert — it is a reservoir of RAGE ligands that continuously activates residual HSCs (Eq. 3), suppresses HSC apoptosis via NF-*κ*B/Bcl-2, and depletes sRAGE through ongoing hepatocyte damage (Eq. 4). This creates “active irreversibility”: *the scar produces more scar*.

The simulation data confirm this distinction. During F3 abstinence, permanent collagen fraction remains at 0.150 (5× higher than F2 abstinence at 0.033). This reservoir sustains low-level RAGE→HSC→collagen cycling even without exogenous TGF-*β* drive. The system reaches a *fibrotic equilibrium* — not progression, not resolution, but self-sustaining maintenance of the fibrogenic program.

This distinction is the central contribution of the present work: **irreversibility is not just structural resistance to degradation — it is active fibrogenic maintenance mediated by the RAGE perpetuation loop**.

### 7.2 sRAGE as a Reversibility Biomarker

The sRAGE divergence at F2–F3 (Table 3) predicts that serum sRAGE trajectories during fibrosis regression could serve as a dynamic biomarker of the reversibility boundary. This is independently testable: serial sRAGE measurements in ALD patients undergoing alcohol cessation, stratified by baseline fibrosis stage, should reveal sRAGE recovery in F2 patients but persistent depression in F3+. Existing cross-sectional data showing inverse correlation between sRAGE and fibrosis stage [6, 17] are consistent with this prediction but do not test the dynamic trajectory. The model predicts that *longitudinal* sRAGE monitoring would reveal the moment the RAGE loop engages: the point where sRAGE stops recovering despite injury removal.

### 7.3 Therapeutic Implications: Breaking the Loop

The active irreversibility model identifies four distinct points where the RAGE perpetuation loop could be interrupted:

i. **RAGE receptor blockade**. Small-molecule RAGE antagonists (e.g., FPS-ZM1) block AGE-RAGE binding and have demonstrated anti-fibrotic effects in preclinical renal and cardiac models. Azeliragon (TTP488), an oral RAGE inhibitor developed for Alzheimer’s disease (Phase 3, failed for AD), has not been tested in hepatic fibrosis. Neither compound has entered liver-specific clinical trials. Our model predicts that RAGE blockade at F3 would disengage the perpetuation loop by eliminating the RAGE→HSC activation signal, potentially allowing endogenous resolution mechanisms (NK cell surveillance, Ly6C-lo macrophages, inactivated HSC MMP production) to outpace residual fibrogenesis — even though the crosslinked collagen itself remains structurally intact.
ii. **Exogenous sRAGE administration**. Recombinant sRAGE acts as a decoy receptor, scavenging AGE ligands before they reach membrane RAGE. Preclinical evidence shows anti-fibrotic efficacy in renal fibrosis (attenuated fibronectin, *α*-SMA, and type I collagen via MAPK/NF-*κ*B inhibition) and cardiac fibrosis (reduced collagen deposition and interstitial *α*-SMA+ cells). sRAGE has **not** been tested in hepatic fibrosis models. Our model specifically predicts that exogenous sRAGE would be most effective at F3, where endogenous sRAGE is depleted but sufficient hepatocyte mass remains for regeneration once the RAGE drive is attenuated.
iii. **AGE formation inhibitors**. Aminoguanidine and pyridoxamine prevent new AGE crosslink formation but do not cleave existing crosslinks. These would slow loop escalation during active injury but would not break the loop in established F3+ fibrosis where the AGE reservoir already exists.
iv. **AGE-breakers**. Compounds such as alagebrium (ALT-711) cleave existing AGE crosslinks, potentially eliminating the RAGE ligand reservoir. Alagebrium reached cardiovascular clinical trials but was discontinued in 2009 (Synvista Therapeutics). No AGE-breaker has been tested in hepatic fibrosis. Our model predicts that AGE-breakers would address *both* passive and active irreversibility simultaneously: removing structural crosslinks (restoring MMP susceptibility) while eliminating RAGE ligands (disengaging the perpetuation loop).

#### Stage-dependent therapeutic window

A critical prediction of the model is that all four interventions would exhibit **stage-dependent efficacy**:

- At **F2**: Unnecessary — endogenous resolution succeeds without intervention (∼68% reversal).
- At **F3**: Maximum therapeutic leverage — the RAGE loop is engaged but hepatocyte mass remains sufficient for sRAGE recovery if the loop is broken.
- At **F4**: Reduced efficacy — even with RAGE blockade, insufficient hepatocyte mass may prevent sRAGE recovery, and the extent of structural crosslinking limits MMP-mediated resolution.

This stage-dependent window differs fundamentally from TGF-*β*-targeting strategies (which are most effective during active injury) and may define a complementary therapeutic axis specific to the reversibility boundary.

### 7.4 Limitations and Scope

The model has three ESTIMATED RAGE parameters (Table 1) whose exact values are not experimentally determined. Five parameters are CALIBRATED against multi-objective production validation (HVPG progression, resolution rates, sRAGE divergence), but parameter degeneracy cannot be excluded—alternative parameter combinations may produce similar outputs. The Hill coefficient *n*_2_ = 4 is derived from cascading cooperativities rather than direct measurement. The Gray-Scott substrate pattern for lobule architecture is imposed rather than emergent. Single-lobule simulation does not capture inter-lobular communication.

Several known RAGE-related biological mechanisms are not currently modeled:

- **Alternative RAGE ligands**. The present model considers only AGEs as RAGE activators. HMGB1 released from dying hepatocytes [18] and S100 proteins are additional RAGE ligands that would *strengthen* the feedback loop if incorporated, potentially producing faster loop engagement and lower irreversibility thresholds.
- **BEC–HSC paracrine axis**. RAGE activation on biliary epithelial cells drives BEC proliferation and secretion of Jagged1, which signals neighboring HSCs via Notch/HES1 to promote myofibroblast differentiation — a mechanism particularly relevant to periportal fibrosis and ductular reaction in advanced disease. This intercellular feedback is not captured in the current HSC state model.
- **Epigenetic persistence**. Sustained C/EBP*β* activation from prior alcohol exposure may maintain partial HSC activation independently of ongoing RAGE signaling, providing an additional mechanism for residual fibrogenesis post-abstinence that operates outside our modeled feedback loop.

These omissions represent conservative modeling choices: incorporating any of them would *amplify* the perpetuation loop, likely producing irreversibility at earlier stages. The current model’s predictions may therefore underestimate the true scope of RAGE-mediated active irreversibility.

The model currently addresses alcohol-associated liver disease (ALD). Extension to MASLD is of particular interest because chronic hyperglycemia and insulin resistance — hallmarks of the metabolic syndrome — drive ongoing AGE production via glucose-dependent Maillard chemistry even in the absence of alcohol [19]. The model’s ALD-derived AGE crosslinking rate (*k*_age_ = 0.03) may substantially *underestimate* the glycation burden in diabetic MASLD patients, where a preliminary parameter estimate suggests *k*_age_ ≈ 0.075 (AGE CROSSLINK GLUCOSE = 2.5× ALD baseline). If so, the irreversibility transition would occur at earlier fibrosis stages in MASLD — consistent with clinical observation that metabolic fibrosis can progress rapidly despite the absence of an acute toxin.

## 8 Conclusion

To our knowledge, we present the first computational model of the complete AGE-RAGE-sRAGE feedback loop in hepatic fibrosis. The model produces stage-dependent irreversibility thresholds as an emergent property of the feedback architecture, reproducing clinically observed resolution rates without explicit stage-gating. The sRAGE divergence prediction — recovery at F2 versus persistent depletion at F3 — is independently testable and, if confirmed, would establish sRAGE trajectory as a dynamic biomarker of the fibrosis reversibility boundary.

## Disclosures

### Related Intellectual Property

This work is related to U.S. Provisional Patent Application No. 64/100,396, *“Systems and Methods for Computational Prediction of Hepatic Fibrosis Progression, Irreversibility Detection, and Treatment Management,”* filed June 27, 2026; and U.S. Provisional Patent Application No. 64/101,014, *“Method for Resolution-Invariant Computational Simulation of Biological Tissue Using Variational Lagrangian Discretization,”* filed June 29, 2026. The author has a financial interest in the commercialisation of this technology.

### Use of AI Tools

Portions of this manuscript were drafted with the assistance of large language models (Google Gemini, Anthropic Claude). The author reviewed, verified, and takes full responsibility for all content, including all mathematical derivations, citations, and factual claims.

### Competing Interests

The author declares a financial interest in Decision Sciences, LLC, which may commercialise technologies described herein. Related provisional patent applications are disclosed above.

### Funding

This work was self-funded. No external grants or institutional support were received.

### Data Availability

All simulation parameters are reported in the manuscript (Tables 1–5). Simulation output data supporting the findings are available from the corresponding author upon reasonable request.

### Code Availability

The agent-based simulation engine was developed by the author. Code is available from the corresponding author upon reasonable request, subject to intellectual property restrictions described above.

## Notes

### Competing Interest Statement

The author declares a financial interest in Decision Sciences, LLC, which may commercialize technologies described herein. Related provisional patent applications have been filed (U.S. Provisional Application No. 64/100,396).

